## Supplementary information for "Global regulation via modulation of ribosome pausing by the ABC-F protein EttA"

**Contents:**

Supplementary Figures 1 to 8

Supplementary Tables 1 to 7

Supplementary References

Uncropped gel presented in Figures: 3c-d, 5d and in Supplementary Figures: 1a, 2b and 6c

#### Supplementary Fig. 1

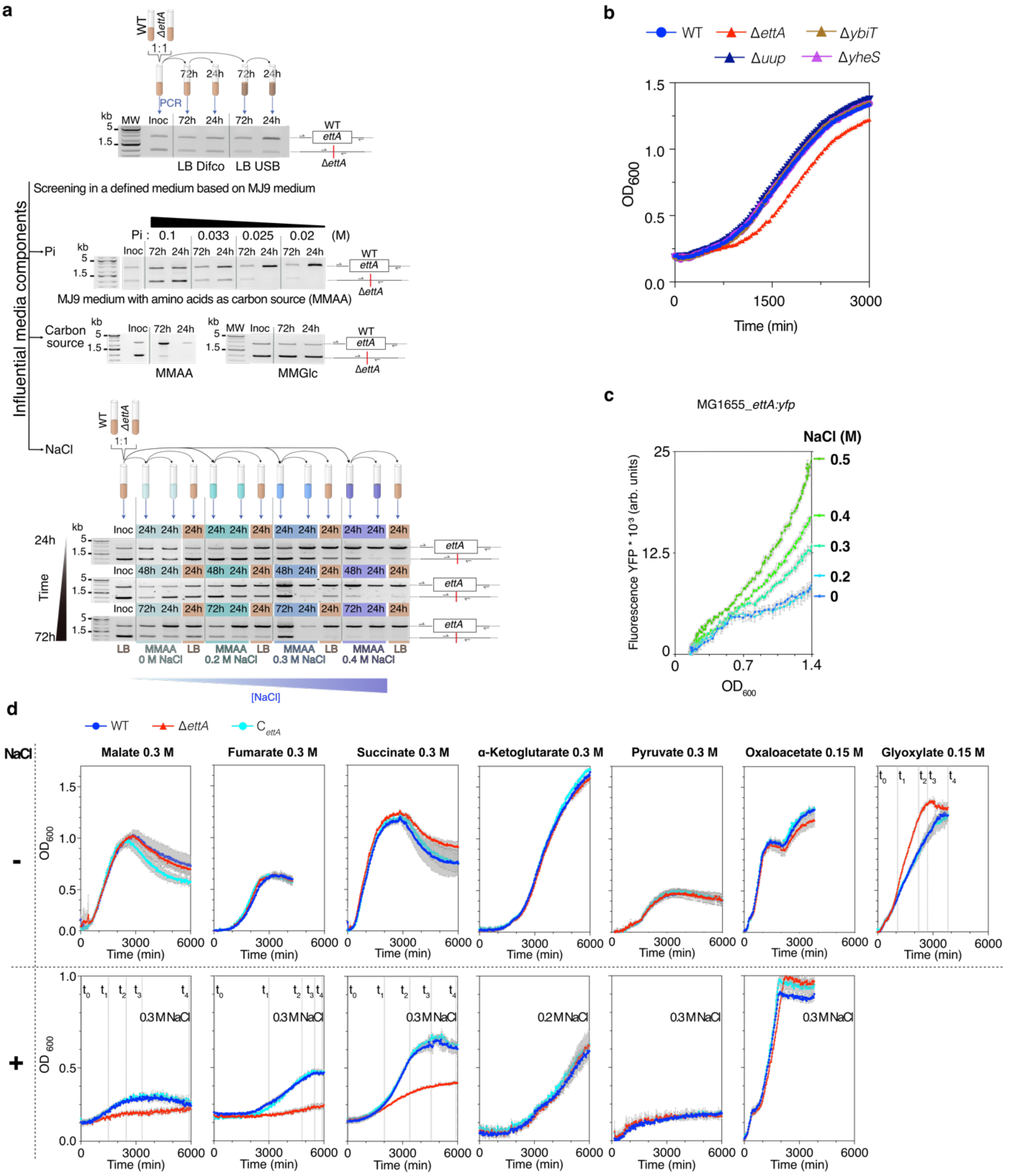


**Supplementary Fig. 1: Phenotypic characterization of culture conditions. a,** Agarose gels showing PCR products from the *ettA* locus in the inoculum, co-culture for 72 h (or 48 h and 24 h in the lower panel) and sub-cultured cultures (+ 24 h) to evaluate the proportion of each strain during fitness experiments between the WT and *ΔettA* strains. Firstly, fitness assay in LB media from two different suppliers: Difco Becton Dickinson and USB-Affymetrix. Secondly from the top, fitness assay in the MJ9 medium[1], with 16 aa as carbon source at 250 mM (glycine at 26.6 mM, alanine at 22.4 mM, valine at 17 mM, leucine at 15.2 mM, isoleucine at 15.2 mM, proline at 17.3 mM, phenylalanine at 12.1 mM, serine at 19 mM, threonine at 16.8 mM, asparagine at 15.1 mM, glutamine 13.7 mM, aspartic acid at 15 mM, glutamic acid at 13.6 mM, arginine at 11.4 mM, histidine at 12.8 mM and lysine at 13.6 mM) where the phosphate concentration has been adjusted to different concentrations (0.1, 0.033, 0.025 and 0.02 M) for 72 hours. Thirdly, fitness in MMAA medium (same as above with 25 mM phosphate) and MMGlc medium where Glucose at 22 mM is used as carbon source in place of the aa. Finally, fitness assays at 24, 48 and 72 hours (from top to bottom) in MMAA medium and in MMAA medium supplemented with increasing concentrations of NaCl (0.2, 0.3 and 0.4 M; left to right). **b**, Graph showing the growth of the WT strain and strains deleted for one of the 4 ABC-F paralogs (Δ*ettA,* Δ*yheS,* Δ*ybiT* and Δ*uup*) in MMAA 0.4 M NaCl culture medium. Error bars represent mean ± standard deviation (s.d.) for triplicate experiments. **c,** Fluorescence of cell cultures expressing *ettA* gene in translational fusion with a *yfp* gene in the MMAA medium in the presence of NaCl at different concentrations (0, 0.2, 0.3, 0.4 and 0.5 M). Curves represent the mean and the standard deviation of the YFP fluorescence at the corresponding $OD_{600}$ during the growth of three independent cultures. **d,** Effect of salt stress on the growth of the WT, Δ*ettA* and $C_{ettA}$ strains in minimum medium (MM) with different intermediate metabolites of the TCA cycle as carbon sources. The $OD_{600}$ was measured every 30 minutes on three independent cultures using a plate reader. Error bars represent mean ± s.d. for triplicate experiments. The time points marked on certain growth curves correspond to the OD values reported in **Fig. 1c**.

#### Supplementary Fig. 2

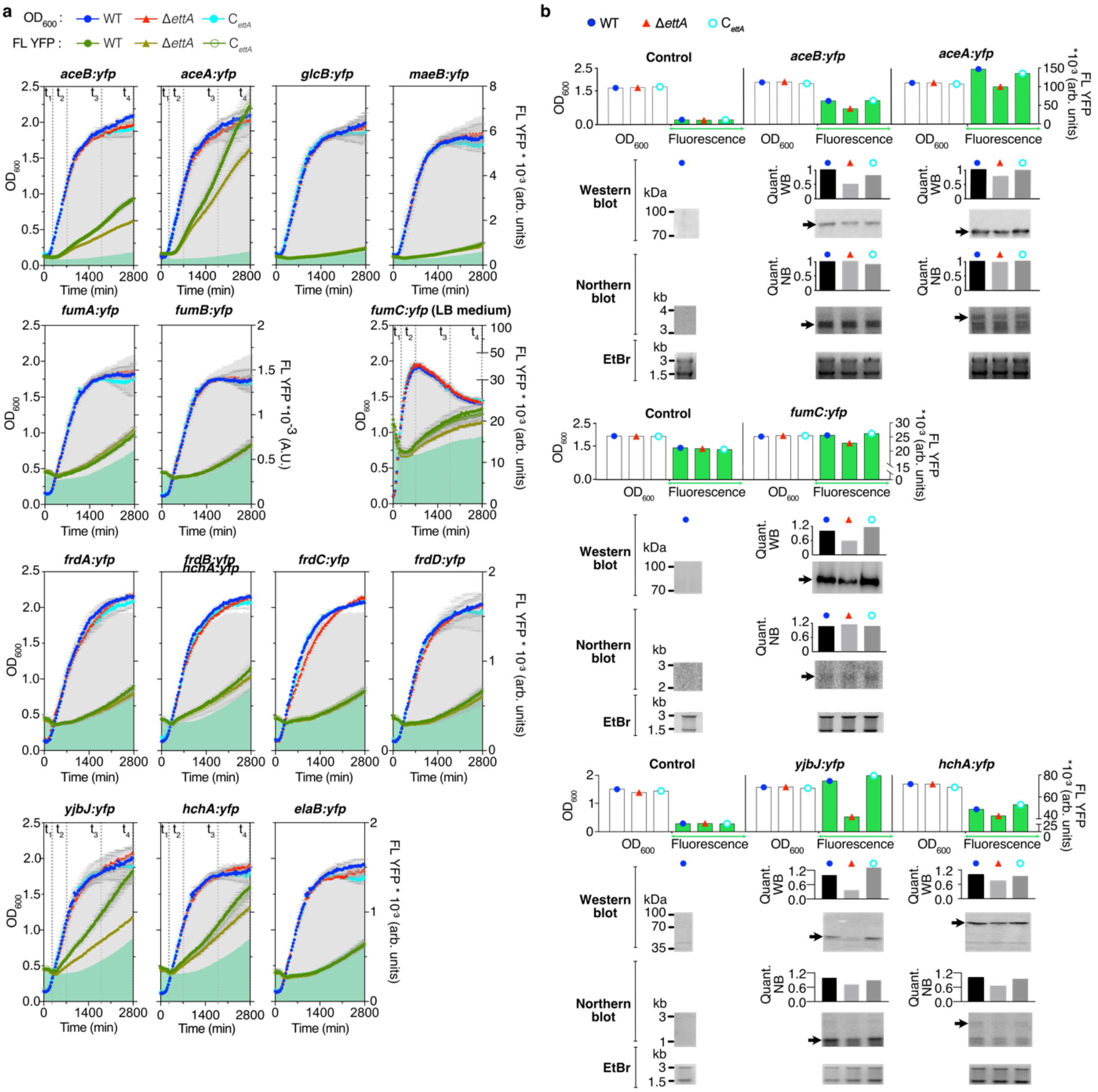


**Supplementary Fig. 2: Validation of the change in expression of some genes in the Δ*ettA* strain with chromosomal *yfp* reporter fusions. a,** Growth curves (blue, cyan and red) and fluorescence emission curves (green gradient) of triplicate cultures of the WT, Δ*ettA* and $C_{ettA}$ strains expressing the *yfp* gene fused to genes involved in the TCA cycle (*aceB:yfp, aceA:yfp, glcB:yfp, maeB:yfp, fumA/B:yfp, frdA/B/C/D:yfp*) and to genes with other functions (*yjbJ:yfp, hchA:yfp* and *elaB:yfp*). Growth was in MMAA medium except for *fumC:yfp* in LB medium. $OD_{600}$ and fluorescence measurements were made using a ClarioStar microplate reader every 30 min. The gray cloud in the graph corresponds to the growth of the WT control strain without a fusion under the same culture condition. The green cloud in the graph corresponds to the residual fluorescence of the WT control strain without YFP fusion. Error bars represent mean ± s.d. for triplicate experiments. The time points marked on certain growth curves correspond to the fluorescence values reported in **Fig. 2f**. **b,** Validation of the EttA-induced deregulation for *aceB/A:yfp*, *fumC:yfp*, *yjbJ:yfp* and *hchA:yfp* genes by western blot and northern blot. Strains with *yfp* fusions to the different targets were grown in MMAA medium except for *fumC:yfp* in LB medium. Top, measurements of the $OD_{600}$ and fluorescence after 48 h of culture in MMAA medium. Control shows background fluorescence from the WT, Δ*ettA* and $C_{ettA}$ strains. Middle, Western-blot with anti-GFP on protein extracts from the same cultures. Bottom, Northern blot on total RNA extracted from the same cultures, detected with a radioactive probe hybridizing to the *yfp* gene. Bands signals were quantified using Fiji

**Supplementary Fig. 2 (cont.)**

software as described in the methods section. Gels stained with Ethidium Bromide are shown at the bottom of the blots as loading control.

#### Supplementary Fig. 3

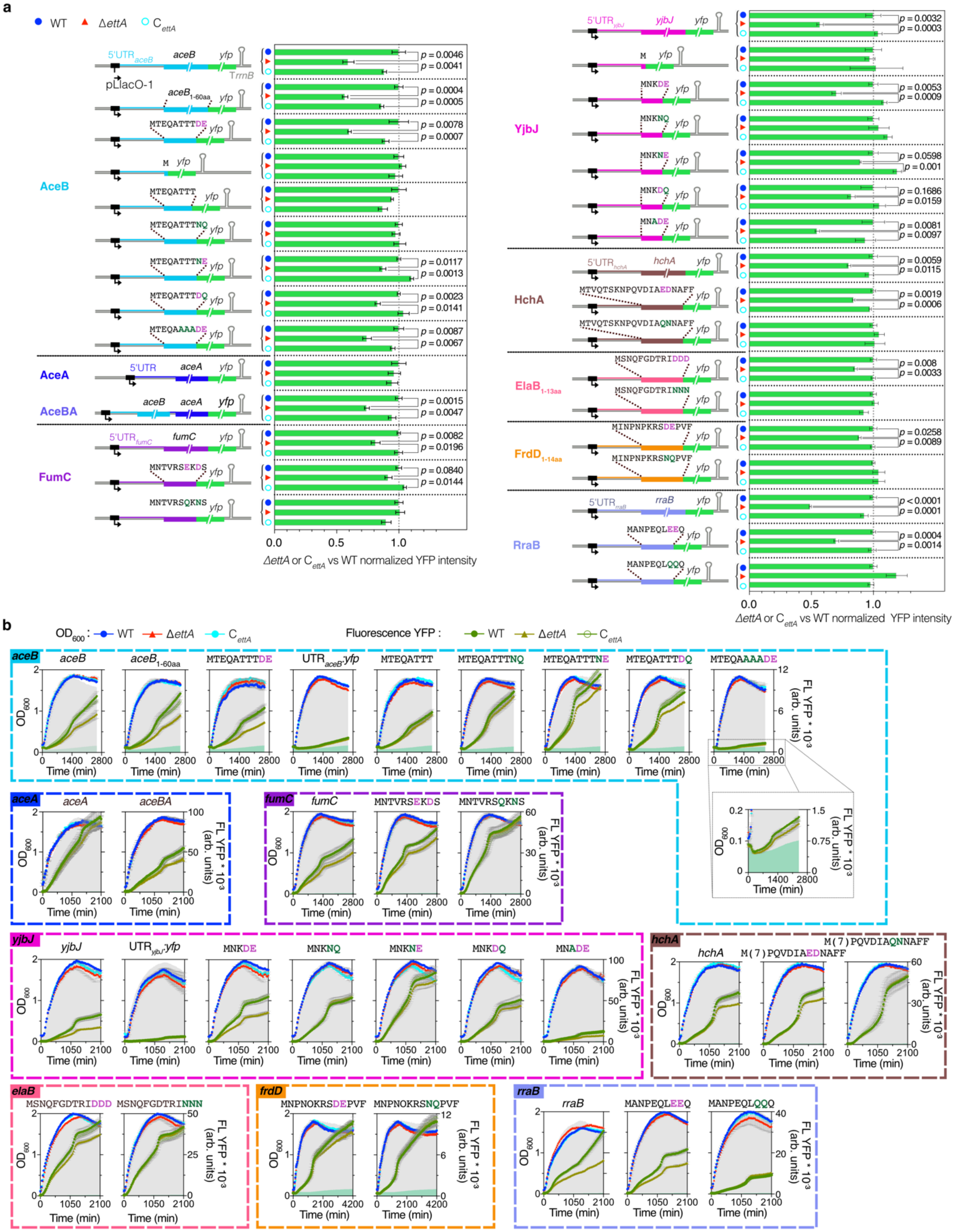


**Supplementary Fig. 3: EttA rescues translation of mRNAs with sequences harboring a motif of two consecutive acidic residues. a,** Histograms showing the ratios of YFP intensity in the WT, Δ*ettA* or $C_{ettA}$ strains normalized over the WT strain (see methods) representing the expression of genes (*aceB, aceA, fumC* and *frdD*, *yjbJ*, *rraB*, *hchA* and *elaB*) in fusion with a *yfp* gene. The various truncated and mutated forms of each fusion tested are shown. All the constructs conserved the 5'UTR sequences of the different genes tested and are expressed from a pMMB plasmid. The tested constructs are shown on the left side of the histograms.

**Supplementary Fig. 3 (cont.)**

Values used to calculate the ratios for all the targets correspond to those at the end of the growth curves presented in panel (**b**). Error bars represent mean ± s.d. for triplicate experiments. For each target, the relative fluorescence value of the WT strain is equal to 1. The *p* values were determined by unpaired two-tailed *t*-tests. **b,** Growth curves of the three strains (WT, Δ*ettA* and $C_{ettA}$) with the different plasmid constructs tested in panel (**a**). Cultures were grown in LB_Amp100 medium in the presence of 1 mM IPTG. $OD_{600}$ and fluorescence measurements were made as described in **Supplementary Fig. 2a.** Error bars represent mean ± s.d. for triplicate experiments. All the growth curves corresponding to different versions of the same target gene are presented within a colored frame corresponding to the color used in panel (**a**): *aceB* (cyan), *aceA* (blue), *fumC* (purple), *yjbJ* (magenta), *hchA* (brown), *elaB* (pink), *frdD* (orange), and *rraB* (light blue).

**Supplementary Fig. 4**

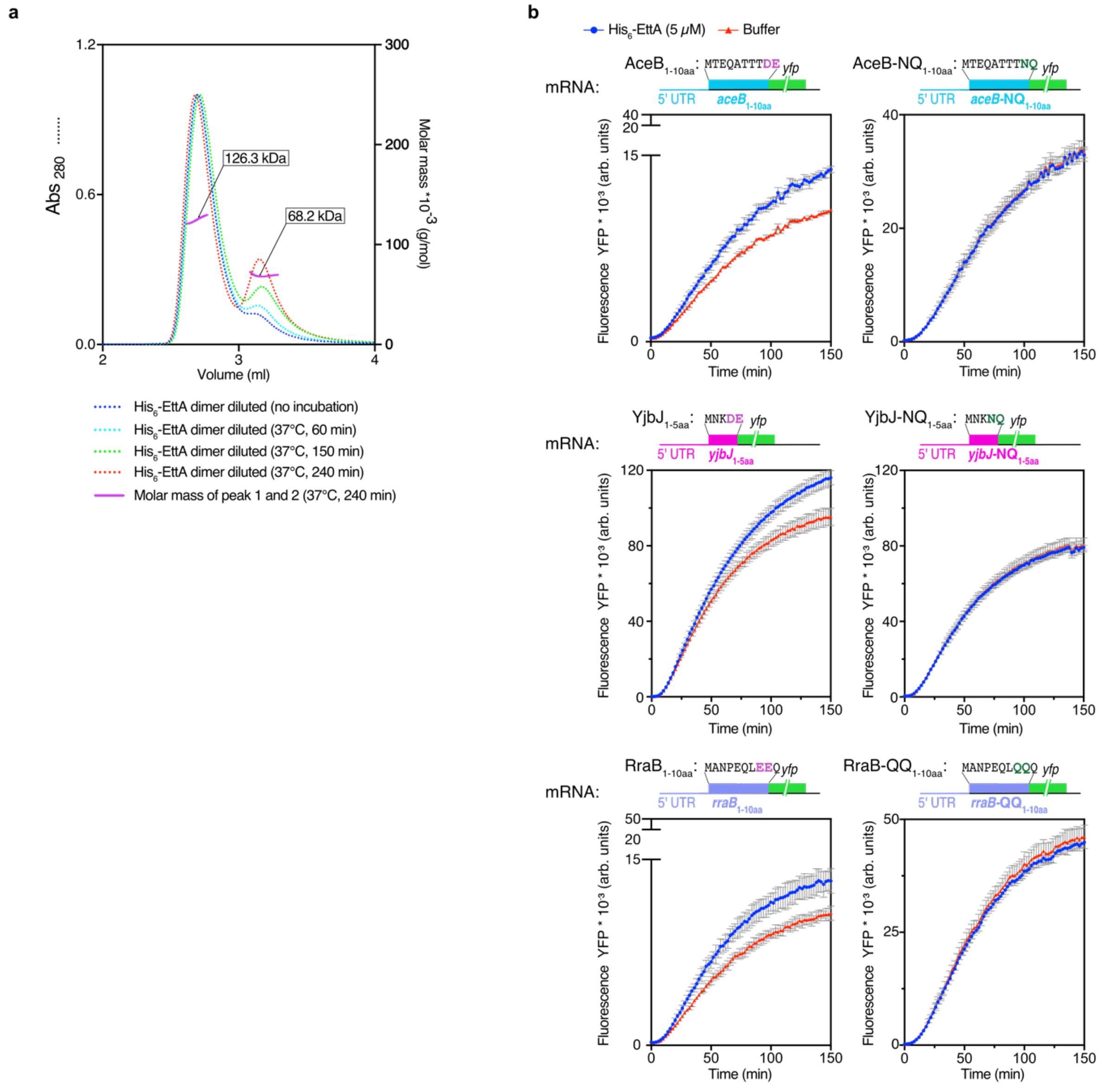


**Supplementary Data Fig. 4:** ***In vitro*** **validation of the involvement of EttA in the regulation of the translation of certain mRNAs. a,** Size exclusion chromatography-multiangle light scattering (SEC-MALS) analysis of the His$_6$-EttA protein purified from *E. coli.* The purified His$_6$-EttA protein at 675 µM was diluted to 40 µM in the same buffer as that used during the purification but with 5% glycerol and heated at 37 °C for 60 min (light blue dotted line), 150 min (green dotted line), or 240 min (red dotted line) before injection. The unheated protein is shown with a blue dotted line. The dotted lines correspond to the absorbance measurements at 280 nm (left axis) and the solid magenta line to the molecular weights (right axis) as a function of the elution volume. The first peak corresponds to His$_6$-EttA protein in its dimeric form (estimated at 126.3 kDa) and the second peak corresponds to the monomeric form (estimated at 68.2 kDa). **b,** Curves showing YFP intensity of *in vitro* translation products from purified mRNAs expressing truncated targets (*aceB*$_{1-10aa}$, *yjbJ*$_{1-5aa}$, *rraB*$_{1-10aa}$) fused with a *yfp* reporter gene, with mutations in acidic residues (NQ or QQ) or without (DE or EE), in the presence (blue) or absence (red) of His$_6$-EttA at 5 µM. Translation reactions were performed with the PURExpress ΔRibosome kit and purified ribosomes from the *E. coli* MRE600 in a final volume of 10 µl for 150 min at 37 °C using a microplate reader.

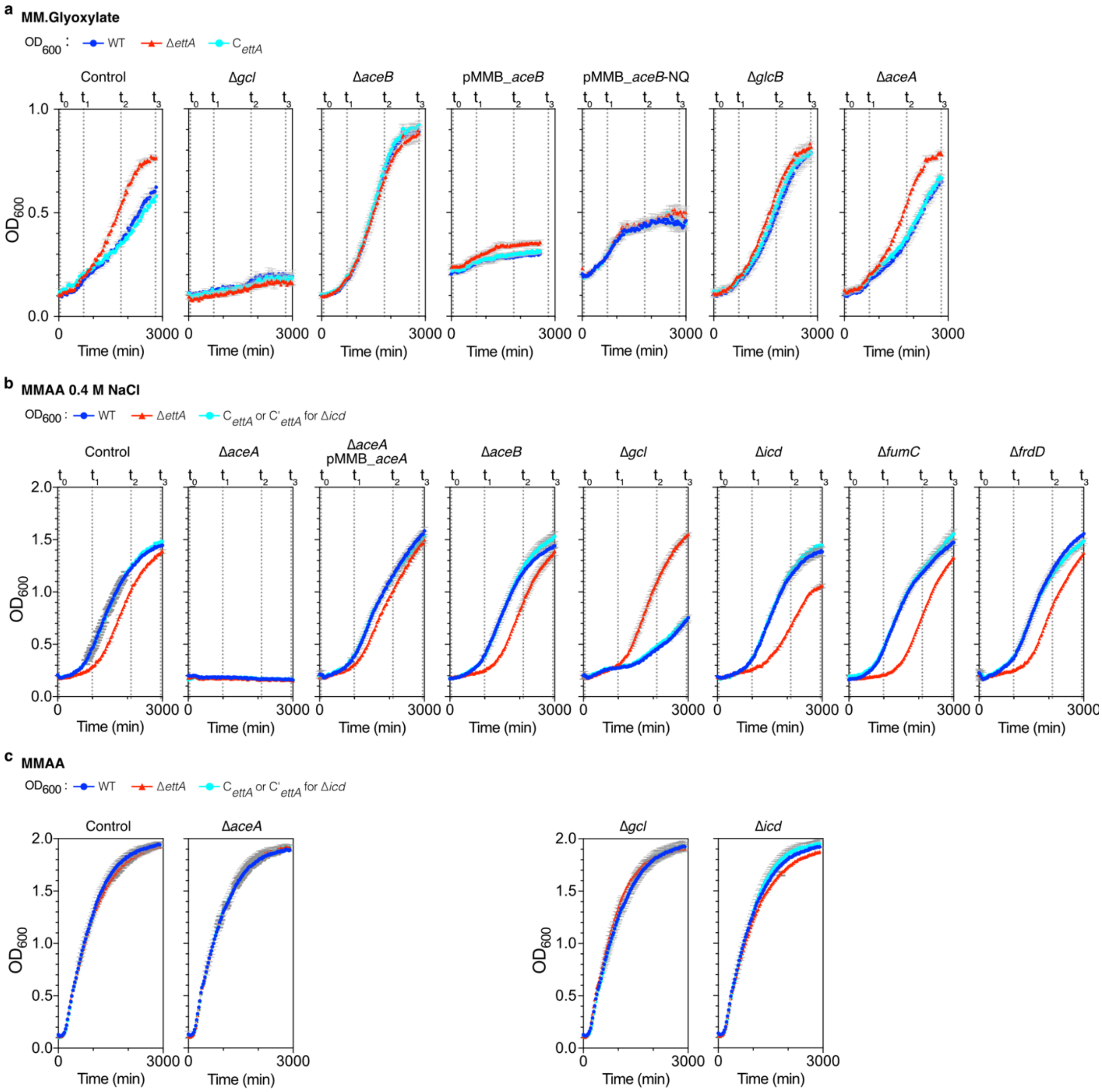


**Supplementary Fig. 5: Growth of the WT, Δ*ettA* and C$_{ettA}$ strains carrying deletions in the genes involved in the metabolism of TCA cycle carbon intermediates in MM.Glyoxylate medium or in MMAA medium in the presence or absence of NaCl. a,** The three strains (WT, Δ*ettA* and C$_{ettA}$) with deletions of the *glc* (Δ*gcl*), *aceB* (Δ*aceB*), *glcB* (Δ*glcB*) or *aceA* (Δ*aceA*) genes or with plasmids over-expressing *aceB* or *aceB*-NQ were grown in MM.Glyoxylate. **b,** The three strains (WT, Δ*ettA* and C$_{ettA}$) with deletions of the *aceB* (Δ*aceB*), *aceA* (Δ*aceA*), *fumC* (Δ*fumC*), *frdD* (Δ*frdD*), *icd* (Δ*icd*) or *gcl* (Δ*gcl*) genes or with a plasmid overexpressing *aceA* were grown in MMAA 0.4 M NaCl **c,** The three strains (WT, Δ*ettA* and C$_{ettA}$) with deletions in the *aceA* (Δ*aceA*), *gcl* (Δ*gcl*) or *icd* (Δ*icd*) genes, were grown in MMAA without NaCl. **a, b and c,** Growth curves were obtained as described in **Supplementary Fig. 1d.** The OD$_{600}$ was measured every 30 minutes on three independent cultures using a plate reader. Error bars represent mean ± s.d. for triplicate experiments. Amp100 and 1 mM IPTG were added to cultures for strains with the pMMB_*aceB*, pMMB_*aceB*-NQ or pMMB_*aceA* plasmids. The time points marked on some growth curves correspond to the OD values reported in **Fig. 4b and c**.

#### Supplementary Fig. 6

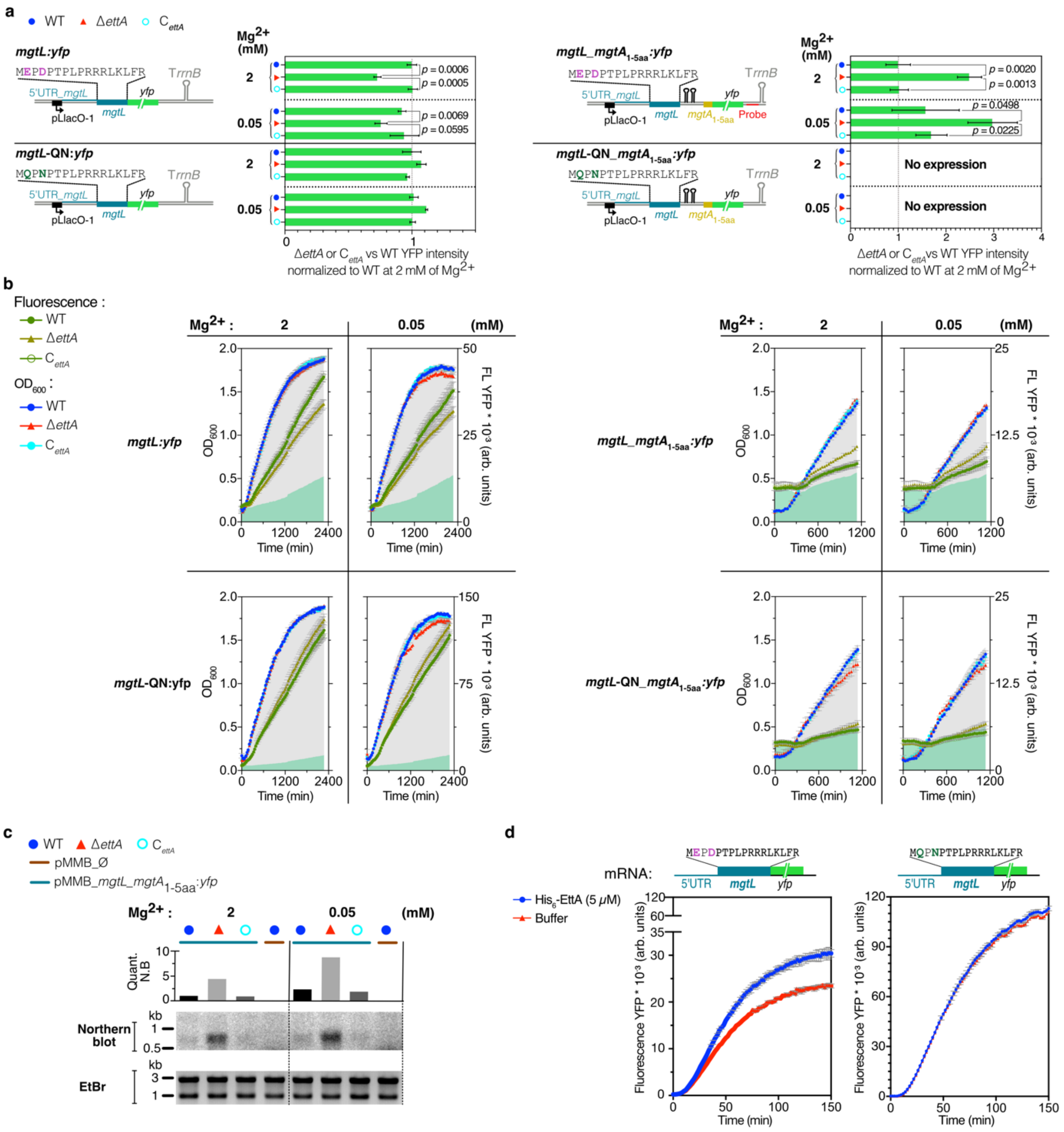


**Supplementary Fig. 6: EttA partly controls the regulation of the leader peptide MgtL. a,** Histograms showing the ratios of YFP intensity in the WT, Δ*ettA* or C$_{ettA}$ strains normalized over the fluorescence of WT strain in the 2 mM Mg$^{2+}$ condition (see Methods), for the *mgtL* (*mgtL:yfp*, left) or for *mgtL* with the 5 first codons of *mgtA* (*mgtL_mgtA*$_{1\text{-}5aa}$*:yfp*, right) fused to a *yfp* gene. The strains were grown in MMAA medium with different concentrations of Mg$^{2+}$ (2 and 0.05 mM) and Amp100. Mutated forms of the same constructs *mgtL*-QN*:yfp* (left, bottom) and *mgtL*-QN_*mgtA*$_{1\text{-}5aa}$*:yfp* (right, bottom) where the Glu2 and Asp4 of MgtL are replaced by Gln and Asn respectively are also shown. All the constructs conserved the 5'UTR sequences of the different genes tested and are expressed on an IPTG-inducible pMMB plasmid. The expression ratios for all cultures are calculated from values at the end of the growth curves presented in panel (**b**). For *mgtL:yfp* and *mgtL*-QN:*yfp, mgtL_mgtA*$_{1\text{-}5aa}$*:yfp* and *mgtL*-QN_*mgtA*$_{1\text{-}5aa}$*:yfp*, the fluorescence value in the WT strain at 2 mM of Mg$^{2+}$ is considered equal to 1 and the values of Δ*ettA* or C$_{ettA}$ strains for each condition are normalized to this value. Ratios are calculated at t=2280 min and t=1140 min for the left and right histograms respectively. All the tested constructs are represented on the left side of the histograms. Error bars correspond to mean ± s.d. for triplicate experiments. The *p* values of unpaired two-tailed *t*-tests are shown on the side of the bars. **b,** Growth curves of the three strains (WT, Δ*ettA* and $C_{ettA}$) with the different constructs shown in panel (**a**). Cultures were performed in MMAA medium at different $Mg^{2+}$ concentrations (2 and 0.05 mM) in the presence of 1 mM IPTG and Amp100. Growth curves were obtained by measuring the $OD_{600}$ every 30 min using a plate reader. $OD_{600}$ and fluorescence measurements were performed as described in **Supplementary Fig. 2a.** Error bars represent mean ± s.d. for triplicate experiments. **c,** Northern blot analysis of *mgtL_mgtA*$_{1\text{-}5aa}$*:yfp* transcript using the oligonucleotide probe for the 3'UTR of the *yfp* (see panel **a**, right). RNAs were extracted after 1140 min of growth from the cultures shown in panel (**b**). Expression of *mgtL_mgtA*$_{1\text{-}5aa}$*:yfp* in the presence of $Mg^{2+}$ at different concentrations (2 and 0.05 mM) was measured for the 3 strains (WT, Δ*ettA* and $C_{ettA}$) and a quantification of the bands is presented in the histogram above the Northern Blot. Staining of the gel with Ethidium Bromide is shown at the bottom of the blot as loading control. **d,** *In vitro* translation of the purified *mgtL:yfp* reporter mRNA in its WT or mutated form (*mgtL*$_{QN}$*:yfp*) and in the presence (blue) or absence (red) of $His_6$-EttA protein at 5 µM as described in **Supplementary Fig. 4b**. Error bars represent mean ± s.d. for triplicate experiments.

#### Supplementary Fig. 7

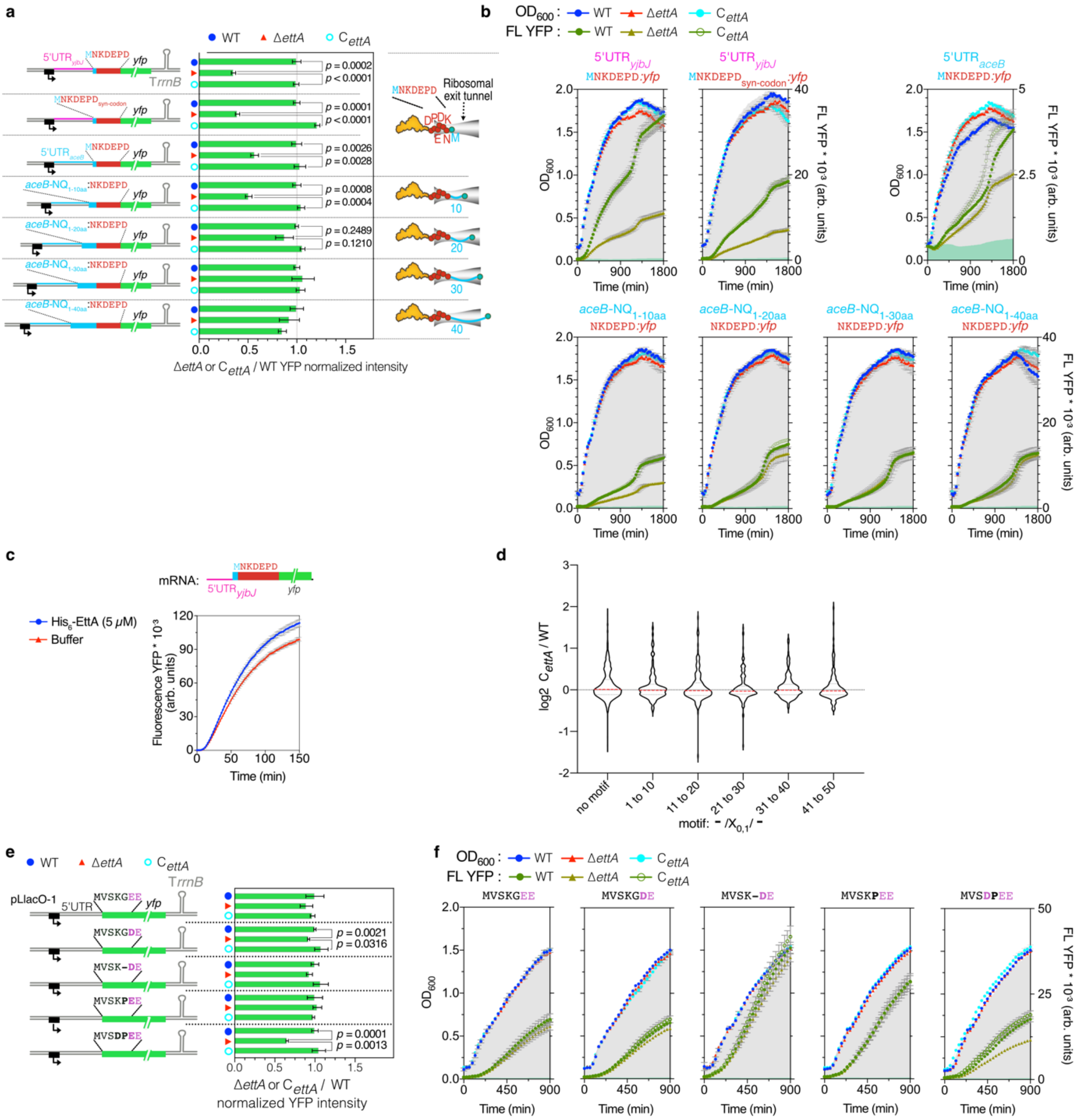


**Supplementary Fig. 7: EttA is a translation factor acting as a facilitator of the synthesis of peptide sequences having the DE or DXE motif in the first 20 amino acids. a,** Histogram showing the translation efficiency of a *yfp* reporter mRNA according to the localization of the NKDEPD motif within the first 40 neosynthesized amino acids, in the three strains (WT, Δ*ettA* and C$_{ettA}$). Translation efficiency is calculated as the ratio of Δ*ettA* / WT or C$_{ettA}$ / WT ratios of YFP intensities. Seven modified constructs were tested: three express the *yfp* gene with a N-terminal MNKDEPD sequence preceded by either the 5'UTR of *aceB* or the 5'UTR of *yjbJ* or the 5'UTR of *yjbJ* followed by a MNKDEPD sequence expressed from an RNA using synonymous aa codons of the MNKDEPD motif (syn-codon, see methods). The other four constructs express the *yfp* gene in fusion with the NKDEPD motif placed either after 10 aa (M(10)NKDEPD), 20 aa (M(20)NKDEPD), 30 aa (M(30)NKDEPD) or 40 aa (M(40)NKDEPD) after the initiation codon, the inserted sequence used for the motif displacement was the *aceB*-NQ. Fluorescence measurements were taken after 1800 min of culture in LB_Amp100 medium with 1 mM IPTG using a microplate reader. Error bars represent mean ± s.d. for triplicate experiments. **b**, Growth curves of the three strains (WT, Δ*ettA* and C$_{ettA}$) with the different plasmid constructs tested in panel (**a**). OD$_{600}$ and fluorescence measurements were carried out as described in **Supplementary Fig. 2a**.

Bacteria were grown in LB_Amp100 medium in the presence of 1 mM IPTG. Error bars represent mean ± s.d. for triplicate experiments. **c,** *In vitro* translation of a *yfp* reporter mRNA fused to a 5'UTR$_{yjbJ}$-MNKDEPD sequence, in the presence (blue) or absence (red) of His$_6$-EttA protein at 5 µM as described in **Supplementary Fig. 4b**. Error bars represent mean ± s.d. for triplicate experiments. **d,** Violin plot showing the expression level of genes containing the E/D-E/D or E/D-X-E/D motif (- /X$_{0,1}$/ -) or not within the first 50 neo-synthesized aa by windows of 10 aa based on the log$_2$ Δ*ettA*/ C$_{ettA}$ ratios obtained in the S$_{15}$ proteomic. The plot show genes expression distributions over *n* = 476 for absence of the motif, *n* = 116 for motif between residues 1 to 10, *n* = 200 for motif between residues 11 to 20, *n* = 176 for motif between residues 21 to 30, *n* = 177 for motif between residues 31 to 40 and *n* = 150 for motif between residues 41 to 50. The red dotted line is the median and the grey ones the quartiles, *p* values were calculated using a two-tailed Mann-Whitney test and were all above 0.05. **e,** Histograms showing the ratios of YFP intensity in Δ*ettA* or C$_{ettA}$ strains vs. WT strain of mutants of the first N-terminal residues of the YFP sequence. The tested constructs are presented on the left side of the histograms. All the constructs have the 5'UTR sequence of the original plasmid pMMBp_EH67-*yfp*. Values used to calculate the ratios for all the constructs correspond to those obtained at t=900 min in the panel (**f**). For each target, the fluorescence value of the WT strain is considered equal to 1. Error bars represent mean ± s.d. for triplicate experiments. **a and e**, The *p* values of unpaired two-tailed *t*-tests are shown on the side of the bars. **f,** Growth curves of the three strains (WT, Δ*ettA* and C$_{ettA}$) with the different plasmid constructs shown in panel (**e**). OD$_{600}$ and fluorescence measurements were performed as described in **Supplementary Fig. 2a**. Error bars represent mean ± s.d. for triplicate experiments. Cultures were performed in LB_Amp100 medium in the presence of 1 mM IPTG.

#### Supplementary Fig. 8

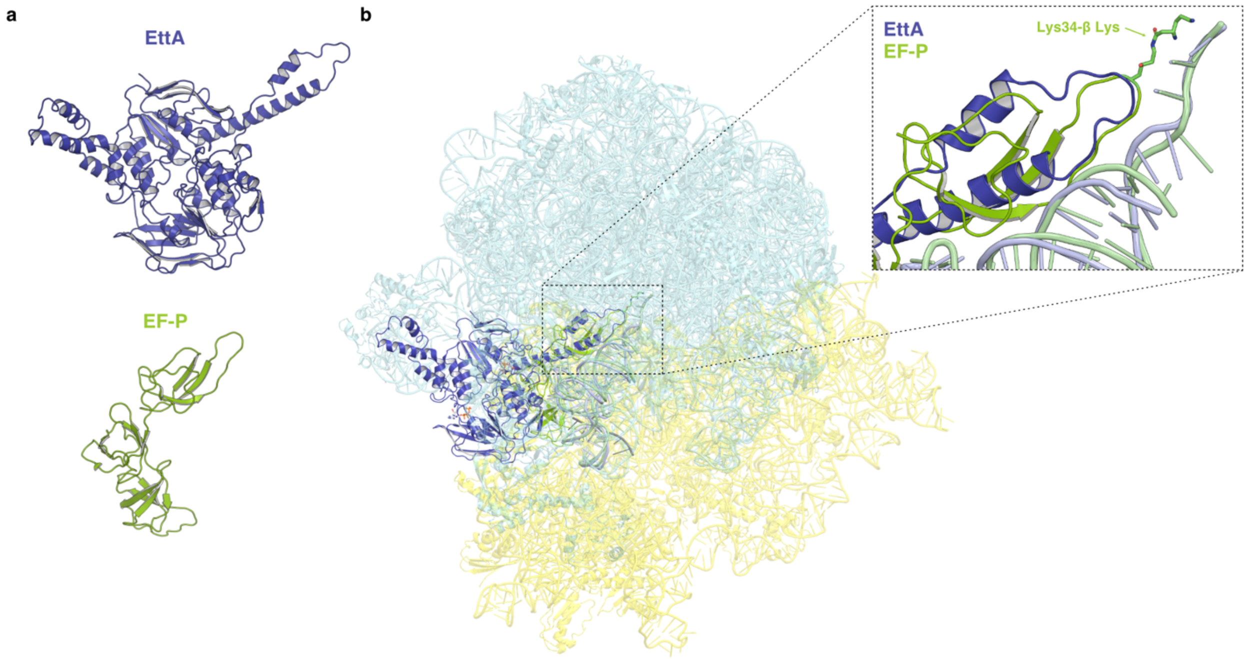


**Supplementary Fig. 8: Structural homology between EttA and EF-P extension toward the PTC.** **a,** Structure of EttA-$EQ_2$ (dark blue) and EF-P isolated from the structure of ribosomal complex (EMD-29398)[2] and (6ENJ)[3] respectively. **b,** Structures of EttA-$EQ_2$ (dark blue) in complex with 70S IC complex (EMD-29398)[2] aligned on the domain V of the 23S rRNA of the structure (6ENJ)[3] of the Polyproline-stalled ribosome in complex with EF-P (green). The 50S (cyan) and 30S (yellow) subunits of the EttA-$EQ_2$ ribosomal complex are presented in transparency, a zoomed view of the extension of EF-P and the extremity of the PtIM of EttA toward the PTC that interact with the CCA acceptor stem of the P-site tRNA is presented, top right.

### Supplementary Tables

#### Supplementary Table 1. Reference strains

| Strains | Genotype/ Parent strain | Method used for strain construction | References |
|---|---|---|---|
| DH5α | F⁻ *endA1 glnV44 thi-1 recA1 relA1 gyrA96 deoR nupG purB20* φ80d*lacZ*ΔM15 Δ(*lacZYA-argF*)U169, hsdR17($r_K^- m_K^+$), λ⁻ | - | [4] |
| MRE600 | F- rna | - | [5] |
| MG1655 | K-12 F⁻ λ⁻ *ilvG⁻ rfb-50 rph-1* | - | [6] |
| Δ*ettA* | MG1655 Δ*ettA* | - | [7] |
| Δ*uup* | MG1655 Δ*uup* | | |
| Δ*ybiT* | MG1655 Δ*ybiT* | | |
| Δ*yheS* | MG1655 Δ*yheS* | | |
| $C_{ettA}$ | MG1655 Δ*ettA, P21:ettA* | pOSIP_CT (P21) plasmid[8] | This study |
| $C'_{ettA}$ | MG1655 Δ*ettA, attB:ettA* | pOSIP_KL (*attB*) plasmid[8] | This study |

#### Supplementary Table 2. Strains with genomic YFP reporter

| Strain name | Method used for strains construction | Selection |
|---|---|---|
| MG1655_*aceA:yfp* | P1 transduction from Venus-YFP (GFP) fusion strains[9] to parental strain MG1655, Δ*ettA* or $C_{ettA}$ | $Cm^R$ |
| Δ*ettA_aceA:yfp* | | |
| $C_{ettA}$_*aceA:yfp* | | |
| MG1655_*aceB:yfp* | | |
| Δ*ettA_aceB:yfp* | | |
| $C_{ettA}$_*aceB:yfp* | | |
| MG1655_*glcB:yfp* | | |
| Δ*ettA_glcB:yfp* | | |
| $C_{ettA}$_*glcB:yfp* | | |
| MG1655_*fumA:yfp* | | |
| Δ*ettA_fumA:yfp* | | |
| $C_{ettA}$_*fumA:yfp* | | |
| MG1655_*fumB:yfp* | | |
| Δ*ettA_fumB:yfp* | | |
| $C_{ettA}$_*fumB:yfp* | | |
| MG1655_*fumC:yfp* | | |
| Δ*ettA_fumC:yfp* | | |
| $C_{ettA}$_*fumC:yfp* | | |
| MG1655_*frdA:yfp* | | |
| Δ*ettA_frdA:yfp* | | |
| $C_{ettA}$_*frdA:yfp* | | |
| MG1655_*frdB:yfp* | | |
| Δ*ettA_frdB:yfp* | | |
| $C_{ettA}$_*frdB:yfp* | | |
| MG1655_*frdC:yfp* | | |
| Δ*ettA_frdC:yfp* | | |
| $C_{ettA}$_*frdC:yfp* | | |
| MG1655_*frdD:yfp* | | |
| Δ*ettA_frdD:yfp* | | |
| $C_{ettA}$_*frdD:yfp* | | |
| MG1655_*elaB:yfp* | | |
| Δ*ettA_elaB:yfp* | | |
| $C_{ettA}$_*elaB:yfp* | | |
| MG1655_*maeB:yfp* | | |
| Δ*ettA_maeB:yfp* | | |
| $C_{ettA}$_*maeB:yfp* | | |
| MG1655_*yjbJ:yfp* | | |
| Δ*ettA_yjbJ:yfp* | The sequence of the *yfp* and $Cm^R$ was PCR amplified from the *aceB:yfp* strain DNA by primers with a 60 nucleotides homology of upstream and downstream of the stop codon of target gene. Genomic integration was performed using lambda-Red recombineering then the construction was transduced into the Δ*ettA* and $C_{ettA}$ strains (See methods). | |
| $C_{ettA}$_*yjbJ:yfp* | | |
| MG1655_*hchA:yfp* | | |
| Δ*ettA_hchA:yfp* | | |
| $C_{ettA}$_*hchA:yfp* | | |

#### Supplementary Table 3. Deletion strains constructed in this study

| Strain name | Method used for strain construction | Selection |
|---|---|---|
| MG1655_Δ*aceA* | P1 transduction from Keio collection deletion strain[10] to parental strain MG1655, Δ*ettA* or $C_{ettA}$ ($C'_{ettA}$ for Δ*icd*) | $Kan^R$ |
| Δ*ettA*_Δ*aceA* | | |
| $C_{ettA}$_Δ*aceA* | | |
| MG1655_Δ*glcB* | | |
| Δ*ettA*_Δ*glcB* | | |
| $C_{ettA}$_Δ*glcB* | | |
| MG1655_Δ*fumC* | | |
| Δ*ettA*_Δ*fumC* | | |
| $C_{ettA}$_Δ*fumC* | | |
| MG1655_Δ*frdD* | | |
| Δ*ettA*_Δ*frdD* | | |
| $C_{ettA}$_Δ*frdD* | | |
| MG1655_Δ*gcl* | | |
| Δ*ettA*_Δ*gcl* | | |
| $C_{ettA}$_Δ*gcl* | | |
| MG1655_Δ*icd* | | |
| Δ*ettA*_Δ*icd* | | |
| $C'_{ettA}$_Δ*icd* | | |

#### Supplementary Table 4. Plasmids constructed during this work

| Plasmids | Selection | References |
|---|---|---|
| pKD46 | Amp$^R$ | 11 |
| pMMBpLacO-1-67EH-*yfp* | Amp$^R$ | 12 |
| pMMB-Ø (pMMB-67EH) | Amp$^R$ | 13 |
| pMMB-*aceB:yfp* | Amp$^R$ | This study |
| pMMB_UTR$_{aceB}$*:yfp* | Amp$^R$ | This study |
| pMMB-*aceB*$_{1-60aa}$*:yfp* | Amp$^R$ | This study |
| pMMB-*aceB*$_{1-10aa}$*:yfp* | Amp$^R$ | This study |
| pMMB-*aceB*$_{1-8aa}$*:yfp* | Amp$^R$ | This study |
| pMMB-*aceB*-NQ$_{1-10aa}$*:yfp* | Amp$^R$ | This study |
| pMMB-*aceB*-D9N$_{1-10aa}$*:yfp* | Amp$^R$ | This study |
| pMMB-*aceB*-E10Q$_{1-10aa}$*:yfp* | Amp$^R$ | This study |
| pMMB-*aceB*-TTT/AAA$_{1-10aa}$*:yfp* | Amp$^R$ | This study |
| pMMB-*aceB*-NQ*:yfp* | Amp$^R$ | This study |
| pMMB-*fumC:yfp* | Amp$^R$ | This study |
| pMMB-*fumC* $_{1-10aa}$*:yfp* | Amp$^R$ | This study |
| pMMB_UTR-*fumC*-QN$_{1-10aa}$*:yfp* | Amp$^R$ | This study |
| pMMB-*rraB:yfp* | Amp$^R$ | This study |
| pMMB-*rraB*$_{1-10aa}$*:yfp* | Amp$^R$ | This study |
| pMMB-*rraB*-QQ$_{1-10aa}$*:yfp* | Amp$^R$ | This study |
| pMMB-*yjbJ:yfp* | Amp$^R$ | This study |
| pMMB-*yjbJ*$_{1-5aa}$*:yfp* | Amp$^R$ | This study |
| pMMB-*yjbJ*-NQ$_{1-5aa}$*:yfp* | Amp$^R$ | This study |
| pMMB-*yjbJ*-D4N$_{1-5aa}$*:yfp* | Amp$^R$ | This study |
| pMMB-*yjbJ*-E5Q$_{1-5aa}$*:yfp* | Amp$^R$ | This study |
| pMMB-*yjbJ*-K3A$_{1-5aa}$*:yfp* | Amp$^R$ | This study |
| pMMB_UTR$_{yjbJ}$*:yfp* | Amp$^R$ | This study |
| pMMB-*hchA:yfp* | Amp$^R$ | This study |
| pMMB-*hchA*$_{1-20aa}$*:yfp* | Amp$^R$ | This study |
| pMMB-*hchA*-QN$_{1-20aa}$*:yfp* | Amp$^R$ | This study |
| pMMB-*frdD*$_{1-14aa}$*:yfp* | Amp$^R$ | This study |
| pMMB-*frdD*-NQ$_{1-14aa}$*:yfp* | Amp$^R$ | This study |
| pMMB-*elaB*$_{1-15aa}$*:yfp* | Amp$^R$ | This study |
| pMMB-*elaB*-NNN$_{1-15aa}$*:yfp* | Amp$^R$ | This study |
| pMMB-*aceA:yfp* | Amp$^R$ | This study |
| pMMB-*aceB-aceA:yfp* | Amp$^R$ | This study |
| pMMB-*aceB* | Amp$^R$ | This study |
| pMMB-*aceB*-NQ | Amp$^R$ | This study |
| pMMB-*aceA* | Amp$^R$ | This study |
| pMMB-*mgtL_mgtA*$_{1-5aa}$*:yfp* | Amp$^R$ | This study |
| pMMB-*mgtL*-QN_*mgtA*$_{1-5aa}$*:yfp* | Amp$^R$ | This study |
| pMMB-*mgtL:yfp* | Amp$^R$ | This study |
| pMMB-*mgtL*-QN*:yfp* | Amp$^R$ | This study |
| pMMB_UTR$_{aceB}$-MNKDEPD*:yfp* | Amp$^R$ | This study |
| pMMB_UTR$_{yjbJ}$-MNKDEPD*:yfp* | Amp$^R$ | This study |
| pMMB_UTR$_{yjbJ}$-MNKDEPD$_{syn-codon}$*:yfp* | Amp$^R$ | This study |
| pMMB_UTR$_{aceB}$_aceB-NQ$_{1-10aa:}$NKDEPD*: yfp* | Amp$^R$ | This study |
| pMMB_UTR$_{aceB}$_aceB-NQ$_{1-20aa:}$NKDEPD*: yfp* | Amp$^R$ | This study |
| pMMB_UTR$_{aceB}$_aceB-NQ$_{1-30aa:}$NKDEPD*: yfp* | Amp$^R$ | This study |
| pMMB_UTR$_{aceB}$_aceB-NQ$_{1-40aa:}$NKDEPD*: yfp* | Amp$^R$ | This study |
| pMMB-*yfp*$_A$ | Amp$^R$ | 14 |
| pMMB_UTR$_{aceB}$_*yfp*-MVSKGDE | Amp$^R$ | This study |
| pMMB_UTR$_{aceB}$_*yfp*-MVSK-DE | Amp$^R$ | This study |
| pMMB_UTR$_{aceB}$_*yfp*-MVSKPDE | Amp$^R$ | This study |
| pMMB_UTR$_{aceB}$_*yfp*-MVSDPEE | Amp$^R$ | This study |
| pBAD-Ø (pBAD/myc-His A) | Amp$^R$ | Invitrogen |
| pBAD-*His*$_6$*-ettA* | Amp$^R$ | 15 |
| pBAD-*His*$_6$*-ettA*_EQ$_2$ | Amp$^R$ | 15 |

#### Supplementary Table 5. Oligonucleotides pairs used for genomic constructions and validation.

| No. | Name | Sequence (5’→3’) | PCR template | Purpose |
|---|---|---|---|---|
| L41 | EttA_448up-R | CGCAAAGAGTTCCAGAACTTGC | Bacteria | Fitness assays |
| L42 | EttA_439dwn-F | GATCATCGGTGATCTGGCGG | | |
| F01 | EttA-pOSIP-P21-F | GAATTCGAGCTCGGTACCCGGCTTTTACTGCG AGGGTGATC | MG1655 genomic DNA | Genomic complementation of *ettA* gene at P21 locus |
| F02 | EttA-pOSIP-P21-R | CATGCATCTCGAGGCATGCCATTTTACGCATTA CTTCGC | | |
| F03 | EttA-pOSIP-attB-F | GAATTCGAGCTCGGTACCCGGCTTTTACTGCG AGGGTGATCG | | Genomic complementation of *ettA* gene at lambda locus (*attB*) |
| F04 | EttA-pOSIP-attB-R | CATGCATCTCGAGGCATGCCATTTTACGCATTA CTTCGC | | |
| F05 | Primer FLIP_F | ATCTGGTGCTGGGTCTGGTG | Strain to validate | Used to verify the integration of the *ettA* gene on the genome |
| F06 | P4 P21(T) | TAGAACTACCACCTGACC | | |
| F07 | P1 P21(T) | ATCGCCTGTATGAACCTG | | |
| F08 | P4 Lambda (L) | TCTGGTCTGGTAGCAATG | | |
| F09 | P3 Lambda (L) | GGGAATTAATTCTTGAAGACG | | |
| F010 | P2 Lambda (L) | ACTTAACGGCTGACATGG | | |
| F011 | P1 Lambda (L) | GGCATCACGGCAATATAC | | |
| E027 | fusion-yjbJ-yfp-F | CAGAAAGATCAGGCAGAAAAAGAGGTCGTGGA TTGGGAAACCCGCAATGAATATCGCTGGACTA GTGCGGCCGCG | Venus yfp (*gfp)* library[9] genomic DNA | Strain constructions with fluorescent *yfp* reporter |
| E028 | fusion-yjbJ-yfp-R | GAAAAAGAAACCGCGTTAAGAGGGCGTACATC CTTGTACACGTCGGGCAGGAGGGATTAATGTA GGCTGGAGCTGCTTCG | | |
| E031 | fusion-hchA-yfp-F | TTTGCAGCGAATGCGTTGGGTAAACTGGCGGC GCAGGAAATGCTGGCAGCTTACGCGGGTACTA GTGCGGCCGCG | | |
| E032 | fusion-hchA-yfp-R | CGTCGTGAGTACTAACGCGGCCGCGATTGATT ATGCGCTTACATTCAAACGTAACAGGGATGTA GGCTGGAGCTGCTTCG | | |
| E029 | fusion-elaB-yfp-F | CAAGGAATTGGTGTGGGCGCGGCCGTTGGGC TGGTACTAGGACTGTTGCTGGCACGCCGTACT AGTGCGGCCGCG | | |
| E030 | fusion-elaB-yfp-R | AAAGCCTCACATTATACGGGGTACTACAAAAAA AATGCAGTACCCCGGTGTAGGGAGGTTTGTAG GCTGGAGCTGCTTCG | | |
| F018 | yjbJ-dwn-F | ACTGGCGCAATGGTACCTGG | Strain to validate | Verification of the integration of the *yfp* fusion and gene deletion |
| F019 | yjbJ-up-R | TTGAAGCACATGGGCTCTGTG | | |
| F024 | hchA-F -200nt | CACTTGCGACGACGTTGCTC | | |
| F025 | hchA-R +200nt | ATTGCCGAATTGCATGGGGG | | |
| F022 | elaB-F -200nt | TCACTGGCCTGATAAGCCTG | | |
| F023 | elaB-R +220nt | TTTGCTGACTGGCCAGCCAG | | |
| D057 | aceB-F | GTTGTGCTGAACGAAAAGAGC | | |
| F081 | aceB-R up | CTTCAATACCCGCTTTCGCC | | |
| D056 | aceA-F | GGAACAGATCACCACTTCCG | | |
| F080 | aceA-R dwn | GTCGGCCTGTTCGAAACG C | | |
| D062 | glcB-F | GGGCGTTTCTGGTTTAACCG | | |
| F083 | glcB-R | TTACAGCAACAGTACGCCGC | | |
| G033 | gcl-F | TAATGTCTGTCGCATCCCGC | | |
| G034 | gcl-R | CGCAGCTTTCTCAAAACGGG | | |
| G041 | icd-F | GCCGCATTATAGCCTAATAACG | | |
| G042 | icd-R | ACCCCAAAACTACCGAGGGG | | |
| D059 | maeB-F | AGTCTGTAGACTCCGGCAG | | Verification of the integration of the *yfp* fusion |
| E052 | YFP_intern-R | GTGGCGGATCTTGAAGTTGG | | |
| D046 | fumA-F | CGCTTTTAACAGGGCAACGG | | |
| E052 | YFP_intern-R | GTGGCGGATCTTGAAGTTGG | | |
| D047 | fumB-F | AAATGCACTTTGCGTGCCGC | | |
| E052 | YFP_intern-R | GTGGCGGATCTTGAAGTTGG | | |
| D048 | fumC-F | ATAAACAGAGCCGCCCTTCG | | |
| E052 | YFP_intern-R | GTGGCGGATCTTGAAGTTGG | | |
| D049 | frdA-F | GAAGATTACTACGCTGCCGC | | |
| E052 | YFP_intern-R | GTGGCGGATCTTGAAGTTGG | | |
| D050 | frdB-F | GCCAGCTAAACGCGTTTACG | | |
| E052 | YFP_intern-R | GTGGCGGATCTTGAAGTTGG | | |
| D052 | frdC-F | AAACACGTCGATCCGGCTG | | |
| E052 | YFP_intern-R | GTGGCGGATCTTGAAGTTGG | | |
| D053 | frdD-F | TGGGACCAGAGCCAATTATC | | |
| E052 | YFP_intern-R | GTGGCGGATCTTGAAGTTGG | | |
| D048 | fumC-F | ATAAACAGAGCCGCCCTTCG | | Verification of the gene deletion |
| J262 | fumC-R | GTGCCGGACGTTGGTGAGTT | | |
| D053 | frdD-F | TGGGACCAGAGCCAATTATC | | |
| J263 | frdD-R | CAAATGTGGAGCAAGAGGCG | | |

#### Supplementary Table 6. Oligonucleotide pairs used for pMMB plasmid constructions.

| No. | Name | Sequence (5'→3') | PCR template | Purpose |
|---|---|---|---|---|
| G029 | start_yfp-F | GTGAGCAAGGGCGAGGAG | pMMBpLlacO-1-67EH-*yfp* | pMMB backbone |
| G072 | pMMB_pLac-R | TGTGCTCAGTATCTTGTTATCCGC | | |
| H009 | utr_aceB_pLac-F | ATAACAAGATACTGAGCACAGTGCTGAACGAAAAGAGCAC | MG1655 genomic DNA | Insert for pMMB-*aceB:yfp* |
| H010 | aceB_pLac-R | AGCTCCTCGCCCTTGCTCACCGCTAACAGGCGGTAGCC | | |
| H023 | 60AA_aceB-yfp-R | GCTCCTCGCCCTTGCTCACGTTATCAATATCTTGCTGCTGCTG | pMMB-*aceB:yfp* | Construction pMMB-*aceB*$_{1\text{-}60aa}$:*yfp* |
| G029 | start_yfp-F | GTGAGCAAGGGCGAGGAG | | |
| H021 | 10AA_aceB-yfp-R | CTCCTCGCCCTTGCTCACTTCATCGGTTGTTGTTGCCTG | pMMB-*aceB:yfp* | Construction pMMB-*aceB*$_{1\text{-}10aa}$:*yfp* |
| G029 | start_yfp-F | GTGAGCAAGGGCGAGGAG | | |
| H054 | 10AA-D9N_E10Q_aceB-yfp-R | CTCGCCCTTGCTCACTTGATTGGTTGTTGTTGCCTGTTC | pMMB-*aceB:yfp* | Construction pMMB-*aceB*-NQ$_{1\text{-}10aa}$: *yfp* |
| G029 | start_yfp-F | GTGAGCAAGGGCGAGGAG | | |
| I063 | 10AA_D9N_aceB-yfp-R | TCCTCGCCCTTGCTCACTTCATTGGTTGTTGTTGCCTGTTCAGTC | pMMB-*aceB*$_{1\text{-}10aa}$:*yfp* | Construction pMMB-*aceB*-D9N$_{1\text{-}10aa}$:*yfp* |
| G029 | start_yfp-F | GTGAGCAAGGGCGAGGAG | | |
| I064 | 10AA_E10Q_aceB-yfp-R | TCCTCGCCCTTGCTCACTTGATCGGTTGTTGTTGCCTGTTCAGTC | pMMB-*aceB*$_{1\text{-}10aa}$:*yfp* | Construction pMMB-*aceB*-E10Q$_{1\text{-}10aa}$: *yfp* |
| G029 | start_yfp-F | GTGAGCAAGGGCGAGGAG | | |
| H055 | 10AA_TTT/AAA_aceB-yfp-R | CTCGCCCTTGCTCACTTCATCGGCTGCTGCTGCCTGTTCAGTCATCGTGC | pMMB-*aceB*$_{1\text{-}10aa}$:*yfp* | Construction pMMB-*aceB*-TTT/AAA $_{1\text{-}10aa}$:*yfp* |
| G029 | start_yfp-F | GTGAGCAAGGGCGAGGAG | | |
| H047 | 8AA_aceB-yfp-R | CTCGCCCTTGCTCACGGTTGTTGTTGCCTGTTCAGTC | pMMB-*aceB*$_{1\text{-}8aa}$:*yfp* | Construction pMMB-*aceB*$_{1\text{-}8aa}$:*yfp* |
| G029 | start_yfp-F | GTGAGCAAGGGCGAGGAG | | |
| H053 | 5AA_aceB-yfp-R | TGAACAGCTCCTCGCCCTTGCTCACTGCCTGTTCAGTCATCGTGC | pMMB-*aceB*$_{1\text{-}5aa}$:*yfp* | Construction pMMB-*aceB*$_{1\text{-}5aa}$:*yfp* |
| G029 | start_yfp-F | GTGAGCAAGGGCGAGGAG | | |
| H034 | utr_aceB-0aa_pLac-R | CTCCTCGCCCTTGCTCATCGTGCAGCTCCTCGTC | pMMB-*aceB:yfp* | Construction pMMB_UTR$_{aceB}$;*yfp* |
| G029 | start_yfp-F | GTGAGCAAGGGCGAGGAG | | |
| I014 | fumC_pMMB-F | ATAACAAGATACTGAGCACAGTGAGCTAAAAGTTGCTTAACGAAAG | MG1655 genomic DNA | Insert for pMMB-*fumC:yfp* |
| I013 | fumC_pMMB-R | AGCTCCTCGCCCTTGCTCACACGCCCGGCTTTCATACTGC | | |
| I016 | 10aa_fumC-R | AGCTCCTCGCCCTTGCTCACCGAATCTTTTTCGCTGCGTACTG | pMMB-*fumC:yfp* | Construction pMMB-*fumC*$_{1\text{-}10}$:*yfp* |
| G029 | start_yfp-F | GTGAGCAAGGGCGAGGAG | | |
| I017 | 10aa_E7Q_D9N_fumC-R | AGCTCCTCGCCCTTGCTCACCGAATTTTTTTGGCTGCGTACTGTATTC | | Construction pMMB-*fumC*-QN$_{1\text{-}10}$aa:*yfp* |
| G029 | start_yfp-F | GTGAGCAAGGGCGAGGAG | | |
| I021 | utr_frdD_pMMB-F | ATAACAAGATACTGAGCACATGGGCGGTAACTGTGGTTGC | MG1655 genomic DNA | Insert for pMMB-*frdD:yfp* |
| I022 | utr_frdD_pMMB-R | AGCTCCTCGCCCTTGCTCACGATTGTAACGACACCAATCAGC | | |
| I075 | 14aa_frdD_pMMB-R | CCTCGCCCTTGCTCACGAATACCGGTTCGTCAGAACGC | pMMB-*frdD:yfp* | Construction pMMB-*frdD*$_{1\text{-}14aa}$:*yfp* |
| G029 | start_yfp-F | GTGAGCAAGGGCGAGGAG | | |
| I076 | 14aa_D10N_E11Q_frdD-R | CTCGCCCTTGCTCACGAATACCGGTTGGTTAGAACGCTTTGGATTTGGATTAAT | | Construction pMMB-*frdD*-NQ$_{1\text{-}14aa}$:*yfp* |
| G029 | start_yfp-F | GTGAGCAAGGGCGAGGAG | | |
| I069 | rraB-pMMB-F | ATAACAAGATACTGAGCACAATCGGCAGGACATTAAGAGGAATG | MG1655 genomic DNA | Insert for pMMB-*rraB:yfp* |
| I070 | rraB-pMMB-R | AGCTCCTCGCCCTTGCTCACGTGGCGAACTCCGTCATCGTC | | |
| I050 | 10aa_rraB-R | TCCTCGCCCTTGCTCACCTGTTCTTCCAGTTGTTCCGGGTTTGC | pMMB-*rraB:yfp* | Construction pMMB-*rraB*$_{1\text{-}10aa}$:*yfp* |
| G029 | start_yfp-F | GTGAGCAAGGGCGAGGAG | | |
| J007 | 10aa_E8Q_E9Q_rraB_pMMB-R | GCCCTTGCTCACCTGTTGTTGCAGTTGTTCCGGGTTTGCCATG | | Construction pMMB-*rraB*-QQ$_{1\text{-}10aa}$:*yfp* |
| G029 | start_yfp-F | GTGAGCAAGGGCGAGGAG | | |
| I071 | utr_yjbJ-F | ATAACAAGATACTGAGCACAGTACTCTCATTCACAACCTAACGATG | MG1655 genomic DNA | Insert for pMMB-*yjbJ:yfp* |
| I072 | utr_yjbJ-R | AGCTCCTCGCCCTTGCTCACCCAGCGATATTCATTGCGGGTTTC | | |
| I080 | 5aa_yjbJ-R | CCTCGCCCTTGCTCACTTCATCTTTATTCATAATCAAGACCTC | pMMB-*yjbJ:yfp* | Construction pMMB-*yjbJ*$_{1\text{-}5aa}$:*yfp* |
| G029 | start_yfp-F | GTGAGCAAGGGCGAGGAG | | |
| I081 | 5aa_D4N_E5Q_yjbJ-R | CCTCGCCCTTGCTCACTTGATTTTTATTCATAATCAAGACCTCATCGTTAG | | Construction pMMB-*yjbJ*-NQ$_{1\text{-}5aa}$:*yfp* |
| G029 | start_yfp-F | GTGAGCAAGGGCGAGGAG | | |

#### Supplementary Table 6 (continued)

| | | | | |
|---|---|---|---|---|
| I079 | 0aa_yjbJ-R | CCTCGCCCTTGCTCACCATAATCAAGACCTCATCGTTAGGTTG | pMMB-*yjbJ:yfp* | Construction pMMB_UTR$_{yjbJ}$:*yfp* |
| G029 | start_yfp-F | GTGAGCAAGGGCGAGGAG | | |
| J031 | 5aa_D4N_yjbJ-R | TTTTATTCATAATCAAGACCTCATCGTTAG | pMMB-*yjbJ*$_{1\text{-}5aa}$:*yfp* | Construction pMMB-*yjbJ*-D4N$_{1\text{-}5aa}$:*yfp* |
| J030 | 5aa_D4N_yjbJ-F | TGATTATGAATAAAAATGAAGTGAGCAAGGGCGAGG | | |
| J068 | 5aa_E5Q_yjbJ-R | CCTCGCCCTTGCTCACTTGATCTTTATTCATAATCAAGACCTCATC | pMMB-*yjbJ*$_{1\text{-}5aa}$:*yfp* | Construction pMMB-*yjbJ*-E5Q$_{1\text{-}5aa}$:*yfp* |
| G029 | start_yfp-F | GTGAGCAAGGGCGAGGAG | | |
| J067 | 5aa_K3A_yjbJ-R | CCTCGCCCTTGCTCACTTCATCTGCATTCATAATCAAGACCTCATCGTTAG | pMMB-*yjbJ*$_{1\text{-}5aa}$:*yfp* | Construction pMMB-*yjbJ*-K3A$_{1\text{-}5aa}$:*yfp* |
| G029 | start_yfp-F | GTGAGCAAGGGCGAGGAG | | |
| I012 | utr_hchA_pLac-F | ATAACAAGATACTGAGCACAAGCTCAGTCGCAAATATAGTGAC | MG1655 genomic DNA | Insert for pMMB-*hchA:yfp* |
| G028 | utr_hchA_pLac-R | GCTCCTCGCCCTTGCTCACACCCGCGTAAGCTGCCAGCA | | |
| I018 | 20aa-hchA-yfp-R | GCTCCTCGCCCTTGCTCACGAAGAATGCATTATCTTCAGCAATATC | pMMB-*hchA:yfp* | Construction pMMB-*hchA*$_{1\text{-}20aa}$:*yfp* |
| G029 | start_yfp-F | GTGAGCAAGGGCGAGGAG | | |
| I019 | 20aa_E13Q_D4N_hchA-R | AGCTCCTCGCCCTTGCTCACGAAGAATGCATTATTTTGAGCAATATCGAC | pMMB-*hchA:yfp* | Construction pMMB-*hchA*-QN$_{1\text{-}20aa}$: *yfp* |
| G029 | start_yfp-F | GTGAGCAAGGGCGAGGAG | | |
| I059 | utr_elaB_pMMB-F | ATAACAAGATACTGAGCACAAGGTTTTACGCAAATGGAGAACGAG | MG1655 genomic DNA | Insert for pMMB-*elaB:yfp* |
| I060 | utr_elaB_pMMB-R | AGCTCCTCGCCCTTGCTCACACGGCGTGCCAGCAACAGTC | | |
| I062 | 15aa_elaB_pMMB-R | TCCTCGCCCTTGCTCACCGTCAGGTCGTCATCGATACGTG | pMMB-*elaB:yfp* | Construction pMMB-*elaB*$_{1\text{-}15aa}$:*yfp* |
| G029 | start_yfp-F | GTGAGCAAGGGCGAGGAG | | |
| J004 | 15aa_3N_elaB-F | AATAACAACCTGACGGTGAGCAAGGGC | pMMB-*elaB*$_{1\text{-}15aa}$:*yfp* | Construction pMMB-*elaB*-NNN$_{1\text{-}15aa}$: *yfp* |
| J005 | 15aa_3N_elaB-R | CTTGCTCACCGTCAGGTTGTTATTGATACGTGTATCACCAAACTGATTAG | | |
| H011 | aceA_pMMB-F | GATAACAAGATACTGAGCACAGTAAACCACCACATAACTATGGAG | MG1655 genomic DNA | Insert for pMMB-*aceA:yfp* |
| H012 | aceA_pMMB-R | GCTCCTCGCCCTTGCTCACGAACTGCGATTCTTCAGTGGAG | | |
| H009 | utr_aceB_pLac-F | ATAACAAGATACTGAGCACAGTGCTGAACGAAAAGAGCAC | MG1655_*aceA:yfp* genomic DNA | Insert for pMMB-*aceBA:yfp* |
| H012 | aceA_pMMB-R | GCTCCTCGCCCTTGCTCACGAACTGCGATTCTTCAGTGGAG | | |
| H013 | aceA_noyfp_pMMB-F | TAATAATTCGAGCTCGGTACCC | pMMB-*aceA:yfp* | pMMB-*aceA* |
| H014 | aceA_noyfp_pMMB-R | GTACCGAGCTCGAATTATTAGAACTGCGATTCTTCAGTGGAG | | |
| H019 | utr_aceB_noyfp_pLac-F | CTACCGCCTGTTAGCGTAATAATTCGAGCTCGGTACCCG | pMMB-*aceB:yfp* | Construction pMMB-*aceB* |
| H020 | utr_aceB_noyfp_pLac-R | TTACGCTAACAGGCGGTAGC | | |
| J036 | D9N_E10Q_aceB-F | AACTGGCTTTCACAAGGCCGTATG | pMMB-*aceB* | Construction pMMB-*aceB*-NQ |
| J037 | D9N_E10Q_aceB-R | GCCTTGTGAAAGCCAGTTGATTGGTTGTTGTTGCCTGTTCAGTCATC | | |
| G071 | pMMB_pLac_yfp-F | CCCTCGTGACCACCCTGGGC | pMMBpLlacO-1-67EH-*yfp* | pMMB backbone for pMMB-*mgtL_mgtA*$_{1\text{-}5aa}$:*yfp* |
| G072 | pMMB_pLac-R | TGTGCTCAGTATCTTGTTATCCGC | | |
| Twist | mgtL_A-5codon_pMMB-pLac | ATAACAAGATACTGAGCACAGAGATGCTACGAATATTATTGGATTCTCCTTATTATTTGCGGCGCTTTTTTCACTTACCGGAGGTTATATGGAACCTGATCCCACGCCTCTCCCTCGACGGAGATTAAAACTTTTCCGGTAAGCCCGTCTTTTCACGGCGTTACCGGATGCGTAAGGCCGTGACGTTTTAACGTCCCTGCTCAGCTTTATTACCTTCAGGTAAGGCTTCGCCACGCCTGAAGACATTTCTGTACTGTTTCAGACAGTGCGGAGGGACTCCTTATGTTTAAAGAAATTTTTTCCAAAATCGTAAAAATCGTGAGCAAGGGCGAGGAGCTGTTCACCGGGGTGGTGCCCATCCTGGTCGAGCTGGACGGCGACGTAAACGGCCACAAGTTCAGCGTGTCCGGCGAGGGCGAGGGCGATGCCACCTACGGCAAGCTGACCCTGAAGCTGATCTGCACCACCGGCAAGCTGCCCGTGCCCTGGCCCACCCTCGTGACCACCCTGGGC | | Insert for pMMB-*mgtL_mgtA*$_{1\text{-}5aa}$:*yfp* |
| G081 | YFPop-F | TCCAAAATCGTAAAAATCGTGAGC | pMMB-*mgtL_mgtA*$_{1\text{-}5aa}$:*yfp* | Construction pMMB-*mgtL:yfp* |
| G080 | MgtL_YFPop-R | ACGATTTTTACGATTTTGGACCGGAAAAGTTTTAATCTCCGTC | | |

#### Supplementary Table 6 (continued)

| | | | | |
|---|---|---|---|---|
| H024 | E2Q_D4N_ mgtL-F | CAACCTAATCCCACGCCTCTC | pMMB-*mgtL:yfp* pMMB-*mgtL_mgtA*$_{1\text{-}5aa}$*:yfp* | Construction pMMB-*mgtL-QN:yfp* pMMB-*mgtL-QN_mgtA*$_{1\text{-}5aa}$*:yfp* |
| H025 | E2Q_D4N_mgtL-R | AGAGGCGTGGGATTAGGTTGCATAT AACCTCCGGTAAGTGAAAAAAG | | |
| J036 | D9N_E10Q_aceB-F | AACTGGCTTTCACAAGGCCGTATG | pMMB-*aceB:yfp* | Construction pMMB-*aceB*-NQ*:yfp* |
| J037 | D9N_E10Q_aceB-R | GCCTTGTGAAAGCCAGTTGATTGGT TGTTGTTGCCTGTTCAGTCATC | | |
| J038 | NKDEPDyfp-F | AATAAAGATGAACCAGATGTGAGCA AGGGCGAGGAGC | pMMB-*aceB*-NQ*:yfp* | Construction pMMB_UTR$_{aceB}$_MNKDEPD *:yfp* |
| J039 | 0aa_MNKDEPDyfp-R | CTGGTTCATCTTTATTCATCGTGCAG CTCCTCGTCATG | | |
| J038 | NKDEPDyfp-F | AATAAAGATGAACCAGATGTGAGCA AGGGCGAGGAGC | pMMB-*aceB*-NQ*:yfp* | Construction pMMB_ *aceB*-NQ$_{1\text{-}10aa}$ :NKDEPD:*yfp* |
| J040 | 10aa_NKDEPDyfp-R | CTGGTTCATCTTTATTTTGATTGGTT GTTGTTGCCTGTTCAGTCATC | | |
| J038 | NKDEPDyfp-F | AATAAAGATGAACCAGATGTGAGCA AGGGCGAGGAGC | pMMB-*aceB*-NQ*:yfp* | Construction pMMB_*aceB*-NQ$_{1\text{-}20aa}$:NKDEPD:*yfp* |
| J041 | 20aa_NKDEPDyfp-R | CTGGTTCATCTTTATTCTGCTCGCCA TACGGCCTTG | | |
| J038 | NKDEDPyfp-F | AATAAAGATGAACCAGATGTGAGCA AGGGCGAGGAGC | pMMB-*aceB*-NQ*:yfp* | Construction pMMB_*aceB*-NQ$_{1\text{-}30aa}$:NKDEPD:*yfp*$_{1\text{-}}$ 30aa |
| J042 | 30aa_NKDEPDyfp-R | CTGGTTCATCTTTATTTACCGCTTCG GCAGTAAGAATTTGC | | |
| J038 | NKDEPDyfp-F | AATAAAGATGAACCAGATGTGAGCA AGGGCGAGGAGC | pMMB-*aceB*-NQ*:yfp* | Construction pMMB_*aceB*-NQ$_{1\text{-}40aa}$:NKDEPD:*yfp* |
| J043 | 40aa_NKDEPDyfp-R | CTGGTTCATCTTTATTAAAATGCGTC ACCAGCTCAGTCAG | | |
| J034 | NKDEPD_yfp-F | AATAAAGATGAACCAGATGTGAGCA AGGGCGAGGAG CTG | pMMB-*yjbJ:yfp* | Construction pMMB_UTR$_{yjbJ}$_MNKDEPD*: yfp* |
| J035 | NKDEPD_yfp-R | CTGGTTCATCTTTATTCATAATCAAG ACCTCATCGTTAGGTTG | | |
| J065 | NKDEPD_syn-codon_yfp-F | CGAGCCAGACGTGAGCAAGGGCGA GGAGC | pMMB_UTR$_{yjbJ}$_MNKDEP D*:yfp* | Construction pMMB_UTR$_{yjbJ}$_MNKDEPD$_{syn\text{-}codon}$*:yfp* |
| J066 | NKDEPD_syn-codon_yfp-R | CTCACGTCTGGCTCGTCTTTATTCAT AATCAAGACCTCATCG | | |
| J060 | Mutation_motif_in_yfp-F | CTGTTCACCGGGGTGGTGCC | pMMB_*yfp*$_{A}$ (*yfp* optimized) | Construction pMMB_UTR$_{aceB}$_MVSKGDE *:yfp* |
| J061 | motif_MVSKGDE_yfp-R | CACCCCGGTGAACAGTTCATCTCCT TTACTTACCATATTCTGTTT C | | |
| J060 | Mutation_motif_in_yfp-F | CTGTTCACCGGGGTGGTGCC | | Construction pMMB_UTR$_{aceB}$_MVSK-DE*:yfp* |
| J062 | motif_MVSKDE_in_yfp-R | CACCCCGGTGAACAGTTCACTTTAC TTACCATATTCTGTTT CCTGTG | | |
| J060 | Mutation_motif_in_yfp-F | CTGTTCACCGGGGTGGTGCC | | Construction pMMB_UTR$_{aceB}$– MVSKPDE*:yfp* |
| J063 | motif_MVSKPDE_in_yfp-R | CACCCCGGTGAACAGTTCTTCTGGT TTACTTACCATATTCTGTTTCCTG | | |
| J060 | Mutation_motif_in_yfp-F | CTGTTCACCGGGGTGGTGCC | | Construction pMMB_UTR$_{aceB}$_MVSDPEE *:yfp* |
| J064 | motif_MVSDPEE_in_yfp-R | CACCCCGGTGAACAGTTCTTCTGGA TCACTTACCATATTCTGTTTCCTGTG | | |

#### Supplementary Table 7 Oligonucleotides used for Northern blot, IVTA and Toeprinting assay.

**Northern blot probes**

| No. | Name | Sequence (5’→3’) | Target |
|---|---|---|---|
| D072 | Probe_yfp_3'_UTR | CAGCCAAGCTTGCATGCCTGCAGGTCGACTCTAGAGGATCCCCGGG | *mgtA-yfp* |
| C036 | Probe_yfp | CTGGACGTAGCCTTCGGGCATGGCGGACTTGAAGAAGTCGTGCTG | All the other *yfp* fusions |

***In vitro* transcription and translation**

| No. | Name | Sequence (5’→3’) | PCR template | Purpose |
|---|---|---|---|---|
| H033 | T7_pMMB-F | GCGAATTAATACGACTCACTATAGGGA<br>GCGGATAACAAGATACTGAGCACA | pMMB-*aceB*$_{1-10aa}$:*yfp*<br>pMMB-aceB-NQ$_{1-10aa}$:*yfp*<br>pMMB-*yjbJ*$_{1-5aa}$:*yfp*<br>pMMB-*yjbJ*-NQ$_{1-5aa}$:*yfp*<br>pMMB-*rraB*$_{1-10aa}$:*yfp*<br>pMMB-*rraB*-QQ$_{1-10aa}$:*yfp*<br>pMMB_*mgtL*:*yfp*<br>pMMB_*mgtL*-QN:*yfp* | Templates for *in vitro* transcription used in IVTA |
| P010 | After_*yfp*_sequence-R | TGCCTGCAGGTCGACT | | |
| H033 | T7_pMMB_F | GCGAATTAATACGACTCACTATAGGGA<br>GCGGATAACAAGATACTGAGCAC | pMMB-*mgtL*_*mgtA*$_{1-5aa}$:*yfp*<br>pMMB_*mgtL*-QN_*mgtA*$_{1-5aa}$:*yfp* | Templates for *in vitro* transcription used in Toeprinting assay |
| G077 | toe_printing_mgtL-R | GGTTATAATGAATTTTGCTTATTCCTG<br>AAGGTAATAAAGCTGAGC | | |
| H033 | T7_pMMB_F | GCGAATTAATACGACTCACTATAGGGA<br>GCGGATAACAAGATACTGAGCAC | pMMB-*yjbJ*$_{1-5aa}$:*yfp* ;<br>pMMB-*yjbJ*-NQ$_{1-5aa}$:*yfp*<br>pMMB-*rraB*$_{1-10aa}$:*yfp*<br>pMMB-*rraB*-QQ$_{1-10aa}$:*yfp*<br>pMMB-*aceB*$_{1-10aa}$:*yfp*<br>pMMB-aceB-NQ$_{1-10aa}$:*yfp* | Templates for *in vitro* transcription used in Toeprinting assay |
| H059 | R_toe_printing_hybridat ion in *yfp* sequence | GGTTATAATGAATTTTGCTTATTCGTC<br>GCCGTCCAGCTCG | | |
| J109 | TP_NV1_R | Cy5-GGTTATAATGAATTTTGCTTATT | mRNA | Toeprinting assay RT |

#### Uncropped gel

Figure 3c

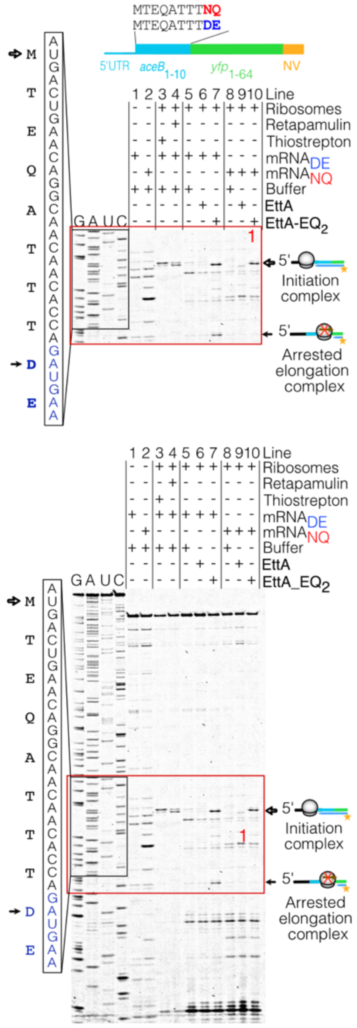


Figure 3d

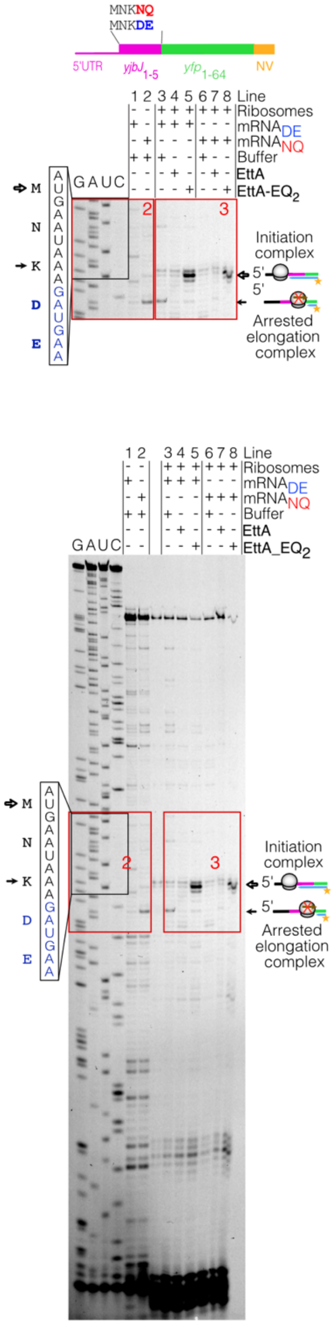


Figure 5d

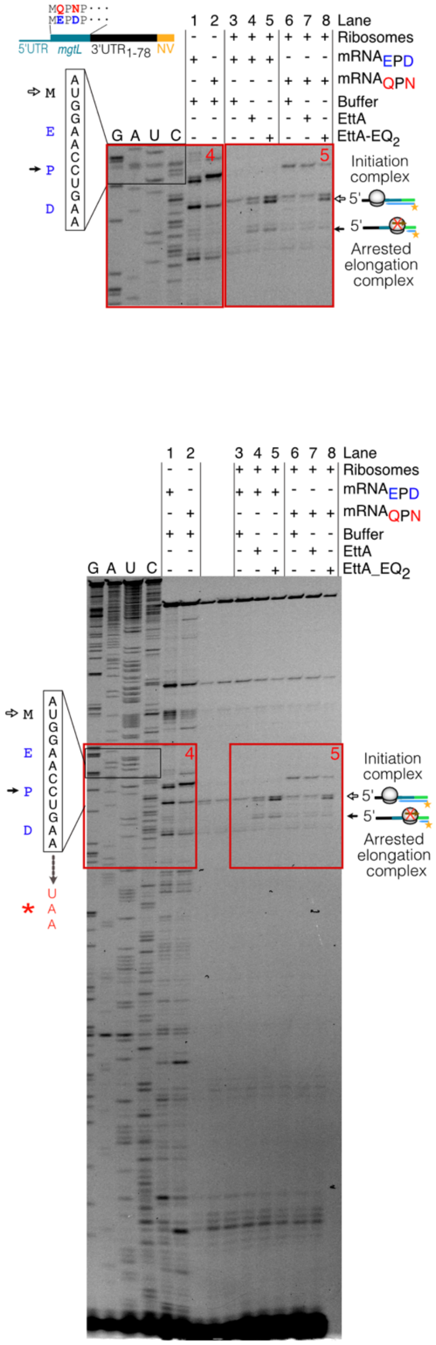

#### Uncropped gel

##### Supplementary Figure 1a

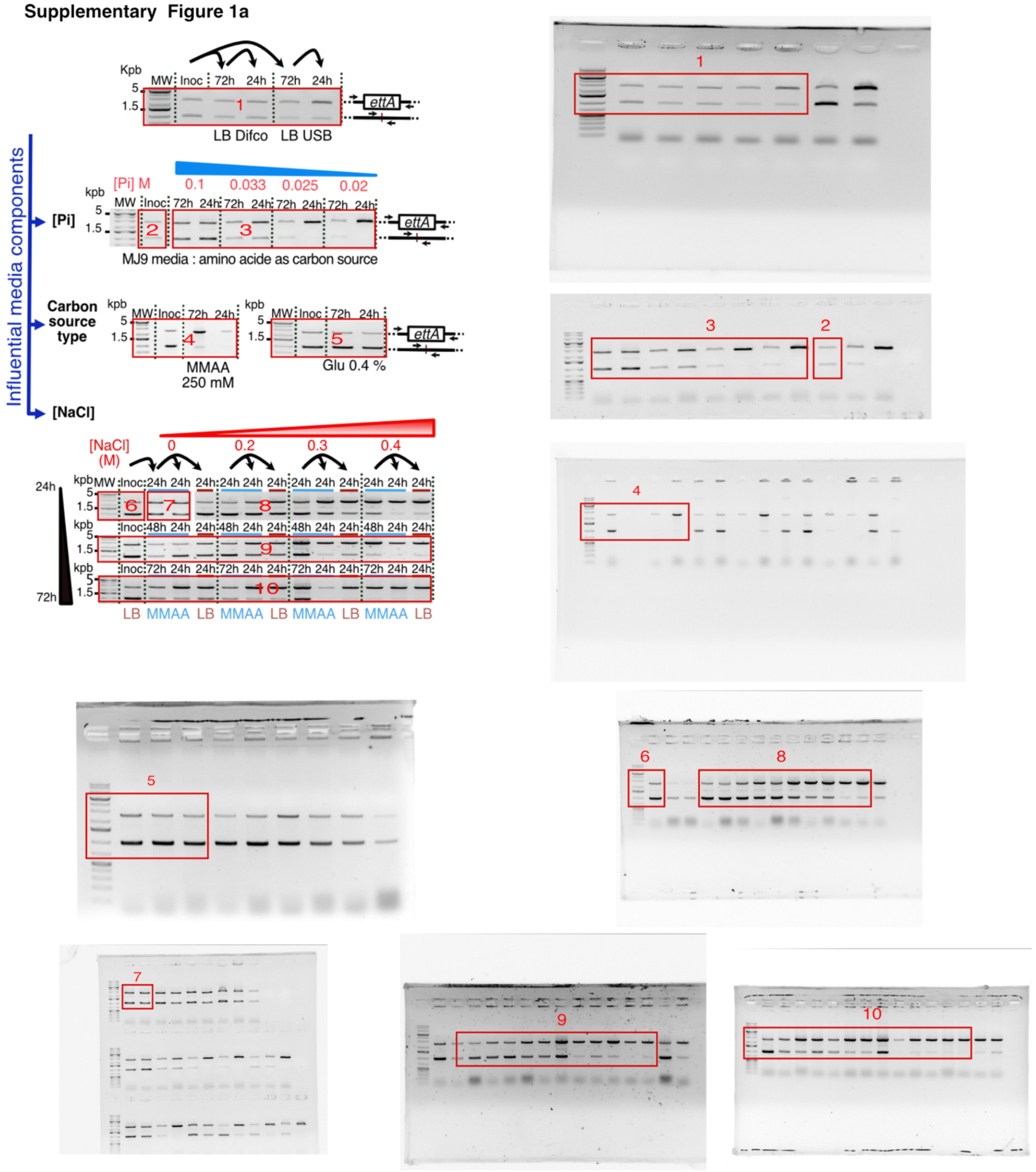

### Uncropped gel

#### Supplementary Figure 2b

**Western Blot :** *aceA:yfp* and *aceB:yfp* (1)

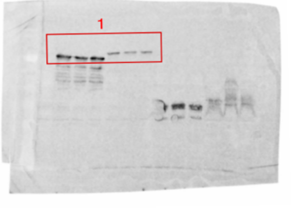


**Northern Blot :** *aceA:yfp* and *aceB:yfp* (2)

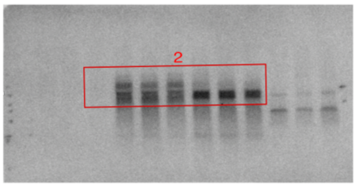


**Western Blot :** *fumC:yfp* (1)

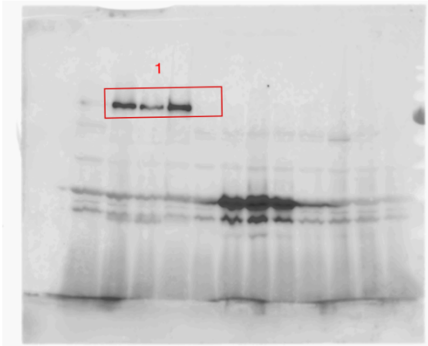


**Northern Blot :** *fumC:yfp* (2)

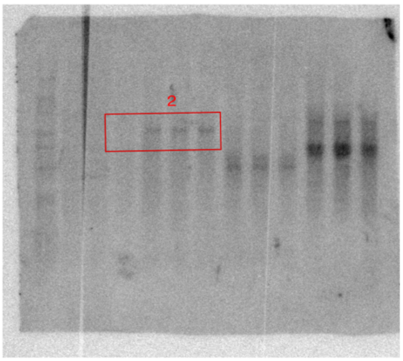


**Western Blot :** *yjbJ:yfp* (1) and *hchA:yfp* (2)

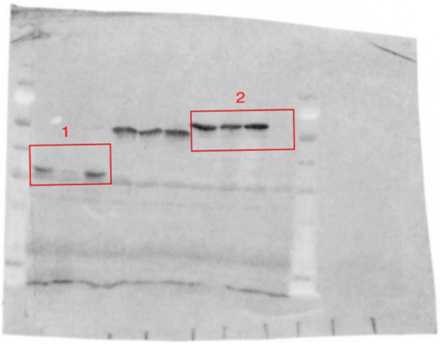


**Northern Blot :** *yjbJ:yfp* (3) and *hchA:yfp* (4)

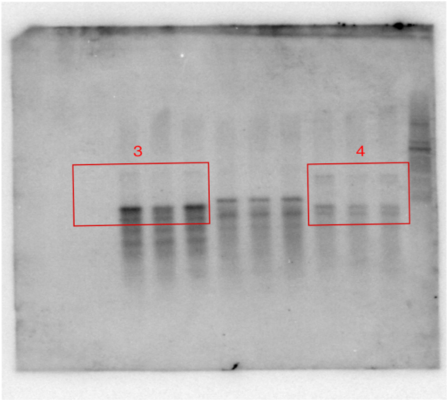


#### Supplementary Figure 6c

**Northern Blot :** *mgtL_mgtA*1-5*:yfp*

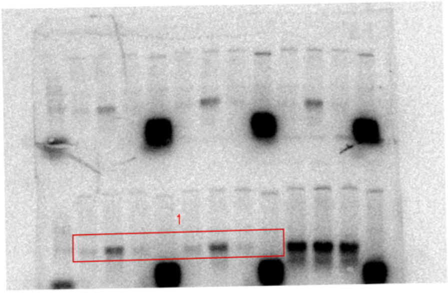
